# tRNA^Ser^ overexpression induces adaptive mutations in NSCLC tumors

**DOI:** 10.1101/2023.05.06.539672

**Authors:** Marta Ferreira, Miguel Pinheiro, Andreia Reis, Ana André, Sara Rocha, Manel A. S. Santos, Mafalda Santos, Carla Oliveira

**Author notes:** Co-senior authors (Mafalda Santos, Carla Oliveira).

## Abstract

tRNAs are a driving force of genome evolution in Yeast and Bacteria. Their deregulation is frequently observed in tumors with Serine tRNAs being often overexpressed. This has important functional consequences, such as increased metabolism and tumor growth. In yeast, time and chemical stimulus boost alterations in the genome driven by tRNA deregulation. Therefore, we hypothesized that tRNA deregulation may contribute to the increased genome instability observed in tumors. To study the effect of tRNA deregulation in tumors, we overexpressed tRNA-Ser-AGA-2-1 in a NSCLC cell line, H460. This cell line and a Mock (control) were xenografted in nude mice and collected at 3 timepoints: T1-Naïve; T2-Treated once with cisplatin/vehicle and; T3) treated twice with cisplatin/vehicle. These tumors were characterized by WES, RNAseq and Mass Spectrometry and the data obtained was integrated. The tumor mutation burden was increased in T3 tRNA^Ser^OE tumors, regardless of treatment. Although in T1 Mock and tRNASer tumors have a similar number of variants, in T2&3, tRNA^Ser^OE tumors display two times more variants than Mock tumors regardless of treatment. Interestingly, tRNA^Ser^OE exclusive variants favor proliferation and therapy resistance, which is in line with the phenotypes observed and supported by RNAseq and proteomics data. In conclusion, tRNA^Ser^OE increases the tumor mutation burden and the variants detected favor tumor growth, proving tRNA deregulation is enough to induce adaptive mutations in the genome.

## Introduction

Genomic instability is a hallmark of cancer (1), which enables the establishment of malignant alterations in a population of cells (2). Both genetic and epigenetic events start accumulating in normal cells long before cancer arises in response to several environmental triggers (3,4). Genomic instability can range from small alterations such as point mutations and microsatellite instability (MSI) to significant structural variations commonly named chromosomal instability (CIN) such as: copy-number variations (CNVs), chromosomal rearrangements, aneuploidy and loss of heterozygosity (LOH) (5). Exposure to carcinogens and faulty DNA repair, replication, cell cycle control or chromosome segregation mechanisms are often the cause of such events (6–9). Besides these highly acknowledged mechanisms, genomic instability has also been reported as a consequence of chronic inflammation (10) or production of genotoxic metabolites by gut microbiota (11,12), highlighting that there may be undisclosed contributors to genomic instability in cancer.

Works in non-mammalian study models, suggest that mutations arising from tRNA deregulation and defective translational machinery, induce important alterations including *de novo* mutations in the genome (13,14). These mutations bestow yeast and bacteria with new adaptive features, allowing them to thrive during stressful events and even develop resistance to therapeutic approaches (15,16). Moreover, a recent paper acknowledges tRNAs as a driving force in genome evolution in yeast (17). tRNA genes are actually capable of modulating gene transcription in a cross-species manner, as they function as DNA insulators from yeast to humans (18–20). Furthermore, some tRNA genes are enriched at topologically associating domains (TADs) and can mediate long-range enhancer-promoter interactions (21). However, the impact of tRNA deregulation in the genome stability of mammalian cells has not been addressed so far. Recently the deregulation of specific tRNAs has been associated with increased aggressiveness in tumor cells (22,23). Overexpression of Serine tRNAs has been detected in breast cancer and considered of prognostic value (24) and it has also been implicated in tumor establishment in an early tumorigenesis lung cancer model (25), In line with this, deregulation of the tRNA pool has been associated to cancer progression, as shown by the distinct tRNA profiles of cancer cell lines and tumors (26). In fact, by increasing the abundance of two tRNAs, Goodarzi et al were able to increase the metastatic potential of human breast cancer cell lines (22).

Given that tRNAs can induce adaptive mutations, which are frequently observed in tumors, and they are also deregulated in cancer, we hypothesized that tRNA deregulation may impact the genome stability of cancer cells. To test that hypothesis, we upregulated a Serine tRNA (tRNA-Ser-AGA-2-1) in a Non-Small Cell Lung Cancer (NSCLC) cell line, H460, and examined its full exome in a longitudinal *in vivo* experiment with or without the exposure to a trigger (cisplatin). Our data shows that the upregulation of tRNASer in these cells increases the tumor mutation burden in the end of the experiment and supports the establishment of adaptive mutations in the genome, unveiling tRNA deregulation as new driver of genomic instability in cancer.

## Results

### Tumor mutation burden is increased in tumor with tRNA^Ser^ upregulation

The ratio of non-synonymous variants per Mb results in a powerful measure: the Tumor mutation burden (TMB) (27). This analysis is particularly relevant in cancers where immunotherapy is an option as is the case of Non-Small Cell Lung Cancer (NSCLC) (28). Tumors with high TMB are designated “hot” tumors and are likely to respond better to this particular therapy (29). Therefore, we calculated the TMB and observed its variation along the experiment in the presence and absence of tRNASer upregulation. Our results show that tumors with upregulation of tRNASer have an increased TMB along the experiment when compared to Mock tumors, regardless of the treatment administered to the mice (Fig. 1A). The differences between Mock and tRNASer tumors in T2 and T3 are exacerbated by cisplatin treatment. However, these results are only significant in T3 tumors treated either with cisplatin or vehicle (Fig. 1A). Additionally, the 1.84-fold increase in TMB observed in T2+Cis, anticipates the results observed in T3 where tRNASer tumors show 1.72-fold increase in TMB when compared to Mock tumors. This observation supports the role of cisplatin as a trigger in our experimental design. Altogether, these results demonstrate that tRNASer tumors are likely hotter tumors and better responders to immunotherapy than their Mock counterparts.

**Figure 1.**
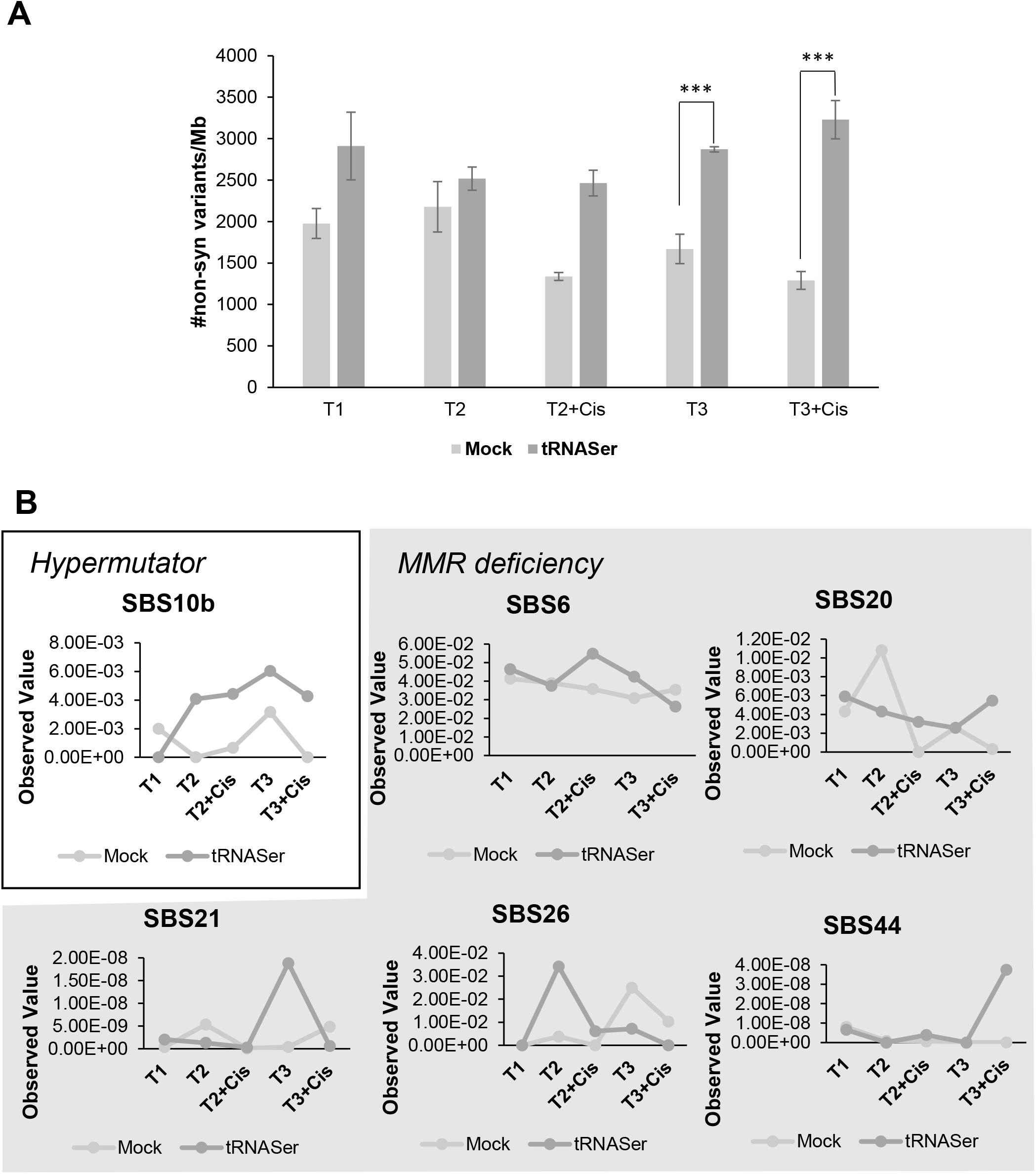
tRNASer tumors have increased TMB and an hypermutator mutational signature. **A)** Quantification of tumor’s Tumor Mutation Burden (TMB) calculated as the total number of non-synonymous variants per MB. tRNASer tumors have increased TMB when compared to Mock tumors from T1 to T3+Cis, but it is only significant in T3 regardless of treatment. Graphic depicts average ± SEM. Data was analyzed with Student’s t-test and significative p-values are shown (***p>0.001). **B)** Stratton signatures variation in tumor cell population along time. tRNASer tumors show an increase in Signature 10b, related to hypermutator phenotype. These tumors, also show increased MMR deficiency signatures in T2 and T3, when compared to their Mock counterparts in the same timepoints. Graphic depicts the observed value of the respective Stratton signatures in tRNASer and Mock tumors along time.

### tRNASer tumors are enriched in hypermutator and MMR deficiency signatures

The somatic mutations in cancer cells arise from multiple processes, which often generate a specific mutational signature composed of base substitutions, small insertions and deletions, genome rearrangements or chromosome copy-number changes (30). After variant calling, we considered the variants that were present in 3 or more tumors as relevant for our analysis, thus decreasing the possibility of considering mutations that arise randomly. The most prominent signatures allowing the distinction of Mock and tRNASer tumors were signatures SBS10b, SBS6, SBS20, SBS21, SBS26 and SBS44 (Full data, Supl. Table 1). Signature SBS10b is associated with a hypermutator phenotype and became predominant in tRNASer tumors from T2 onwards (Fig. 1B). The other signatures identified are attributed to Mismatch Repair (MMR) deficiency and are frequently observed in tumors with microsatellite instability (MSI) (Fig. 1B) (31). These results have a good correlation with the TMB values for these tumors, since tRNASer tumors present an higher number of variants and these tumors have increased hypermutator and MMR deficiency signatures, especially in T2 and T3 tumors either treated or untreated with cisplatin.

### Synonymous and Missense variants are the most common in tRNASer tumors

Next, we predicted the impact of the variants common to 3 or more tumors. In T1, Mock and tRNASer tumors have a very similar number of variants and their predicted impact is also comparable (Fig. 2A). High impact variants are scarce in every timepoint independently of tRNASer expression or treatment status (Fig. 2A). In T2 and T3, tRNASer tumors either treated with vehicle or cisplatin, present a higher number of variants than Mock tumors, which is positively correlated with SBS10b signature increase in these timepoints (Fig.1B, 2A). These variants have mostly predicted Low or Moderate impacts. Regarding the type of variants present in untreated tumors, we observed that after T1, tRNASer tumors show increased numbers of Synonymous variants when compared to Mock tumors, as well as a slight increase in Missense and Intronic variants (Fig. 2B, upper panel). Cisplatin treatment exacerbated these differences, resulting in a clear increase in Missense variants in tRNASer tumors in both T2 and T3 (Fig.2B, lower panel). This increase in Missense mutations, further validates the results from the Stratton Signatures, where the most prevalent mutational signatures were related to MMR deficiency, as well as the significant increased TMB specifically in tRNASer tumors treated with cisplatin in T3 (Fig. 1A).

**Figure 2.**
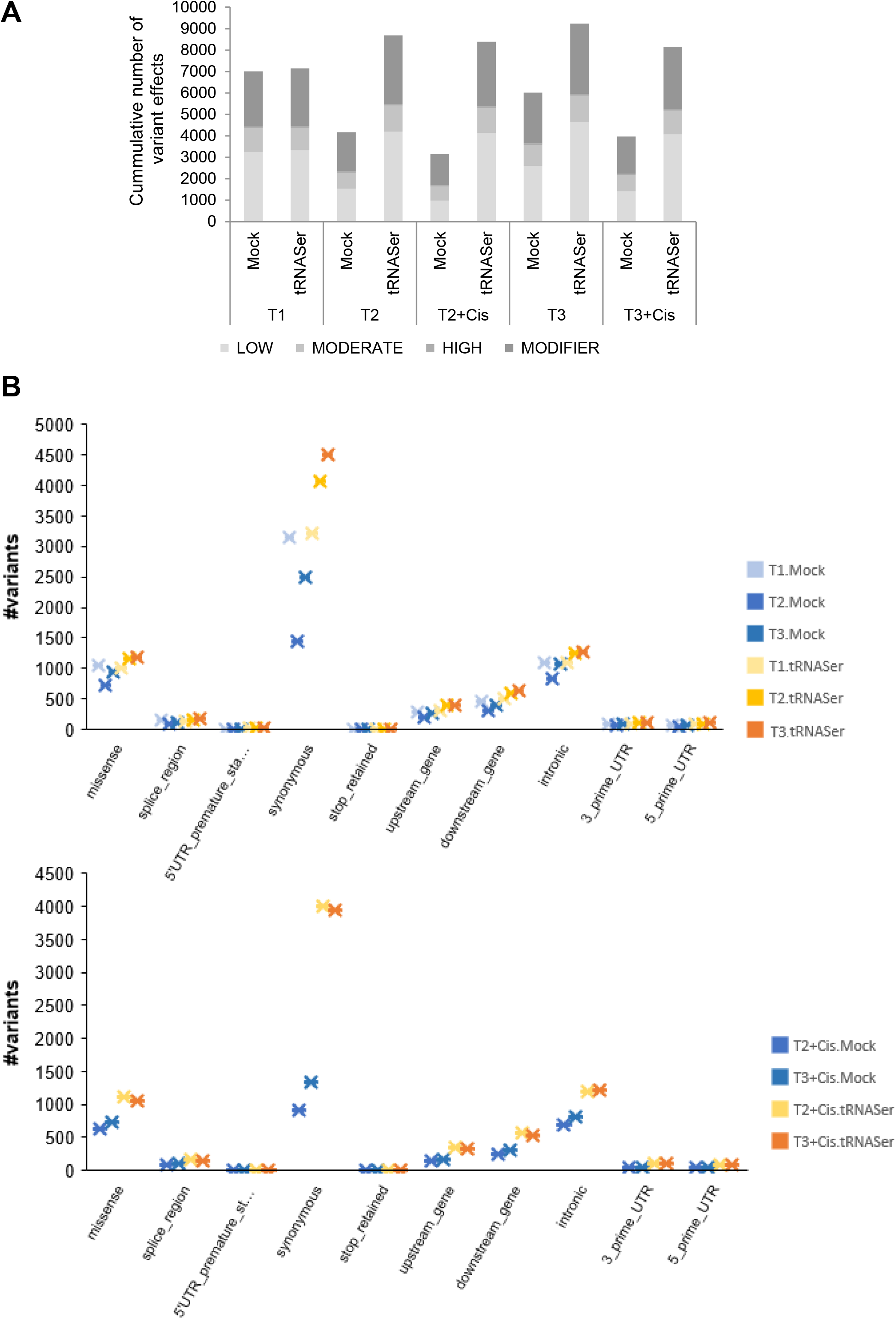
tRNASer tumors display more missense and synonymous variants than Mock tumors. **A)** Graphic depicts the cumulative number of variants present in 3+ tumors according to their predicted impact in tRNASer and Mock tumors along time. In T1, Mock and tRNASer tumors are very similar, but after T2 the total number of variants increases in tRNASer tumors and decreases in Mock tumors. These variants are especially low and moderate impact variants, and only a small number of variants is predicted to have a high impact. **B)** Graphic depicts the number of variants present in 3+ tumors in tRNASer and Mock tumors in the different timepoints. <u>Upper panel:</u> Only variants present in tumors treated with vehicle are represented. In these tumors, we can only observe a difference between tRNASer and Mock tumors regarding synonymous mutations in T2 and T3. <u>Bottom panel:</u> Only variants present in tumors treated with Cisplatin are represented. Cisplatin treatment exacerbates the differences between tRNA and Mock tumors, becoming evident the increase in synonymous, missense and intronic variants in tRNASer tumors.

### Variants detected in tRNASer tumors affect mostly growth and survival genes

Following these results, we characterized the pathways being affected by these variants. Our first observation was that Mock tumors have little or no pathway deregulation upon vehicle or cisplatin treatment, respectively, when compared with tRNASer tumors (Fig. 3). On the other hand, cisplatin treatment of tRNASer tumors increased the number of affected pathways both in T2 and T3. These pathways are predominantly associated with survival and growth, namely: “Signaling by Wnt”, “Signaling by VEGF”, “Regulation of TP53 activity”, “Oncogenic MAPK signaling”, “PI3K-AKT signaling pathway” and “Signaling by PTK6”, among others (Fig. 3).

**Figure 3.**
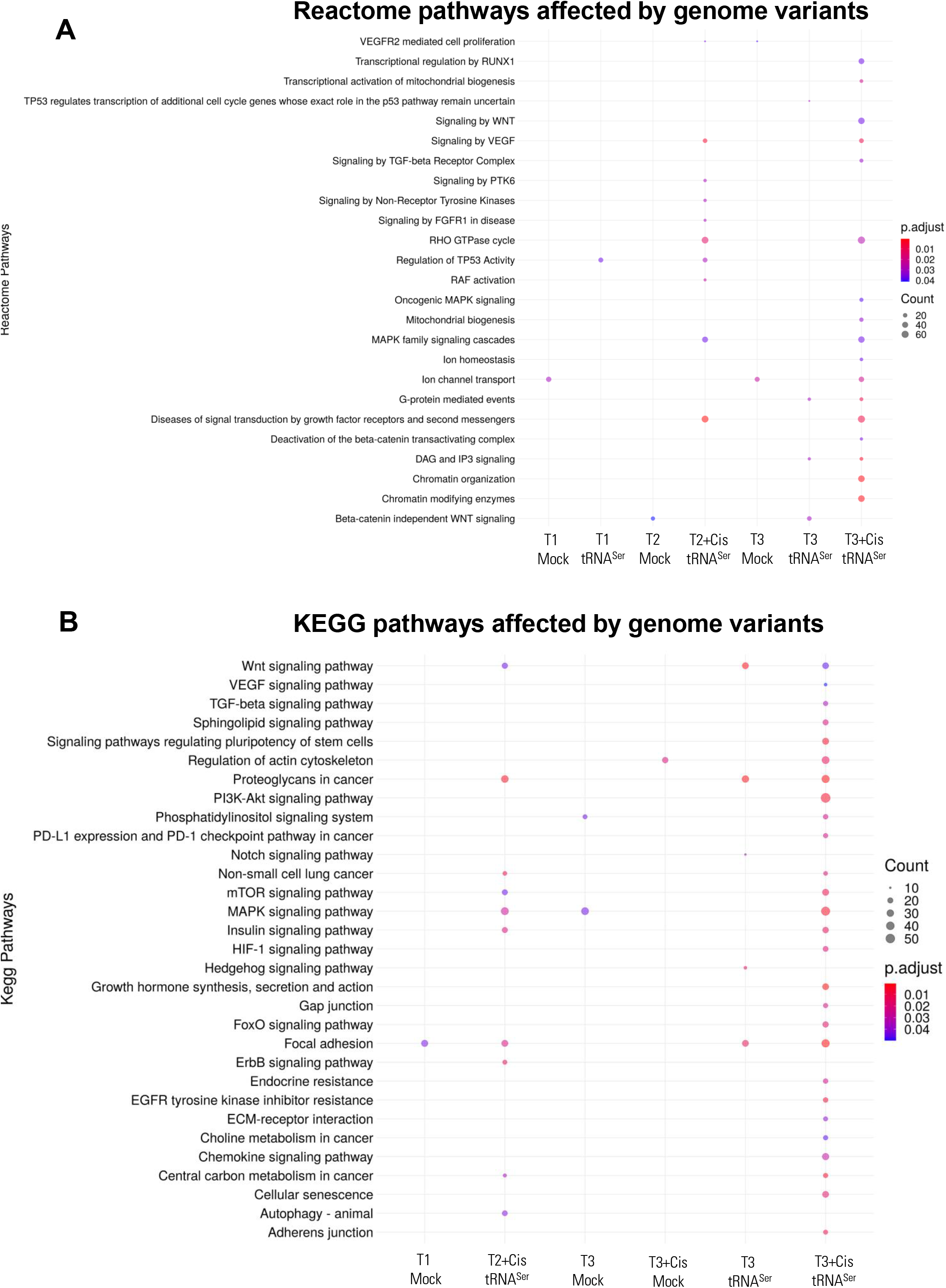
Variants present in tRNASer tumors affect growth and survival signaling pathways. **A)** Reactome pathways affected by variants present in Mock and tRNASer tumors along time. **B)** KEGG pathways affected by variants present in Mock and tRNASer tumors along time. tRNASer tumors have generally a greater number of affected pathways in each time point. Moreover, cisplatin treatment also increases the number of affected pathways. Most of these pathways regulate cell growth and survival. Graphics depict the number of genes (ball size) affected in each pathway and p-value associated (color).

Since the tumors with tRNASer upregulation, do not respond to cisplatin treatment (Fig. S2), this raised the question of whether the genomic variants affecting these pathways were activating them. To answer this question, we resorted to RNAseq and proteomics data. The deregulated GO terms found in RNAseq were represented in circus plots, which shows the up-or downregulated genes associated with each GO term, and calculates the probability of them being activated or repressed in the tumors. Using this strategy, we confirmed the activation of MAPK pathway with GO terms such as “positive regulation of Ras protein signal transduction” and “regulation of MAP kinase activity” being increased in tRNASer tumors in T2+Cis (Fig. 4A) and T3+Cis (Fig. 4B). We also observed an activation of PKB signaling which is a reflection of “PI3K-AKT signaling pathway” and “Signaling by PTK6”. The positive regulation of “Cell Growth” in tRNASer tumors in T2+Cis further supports our hypothesis (Fig. 4A).

**Figure 4.**
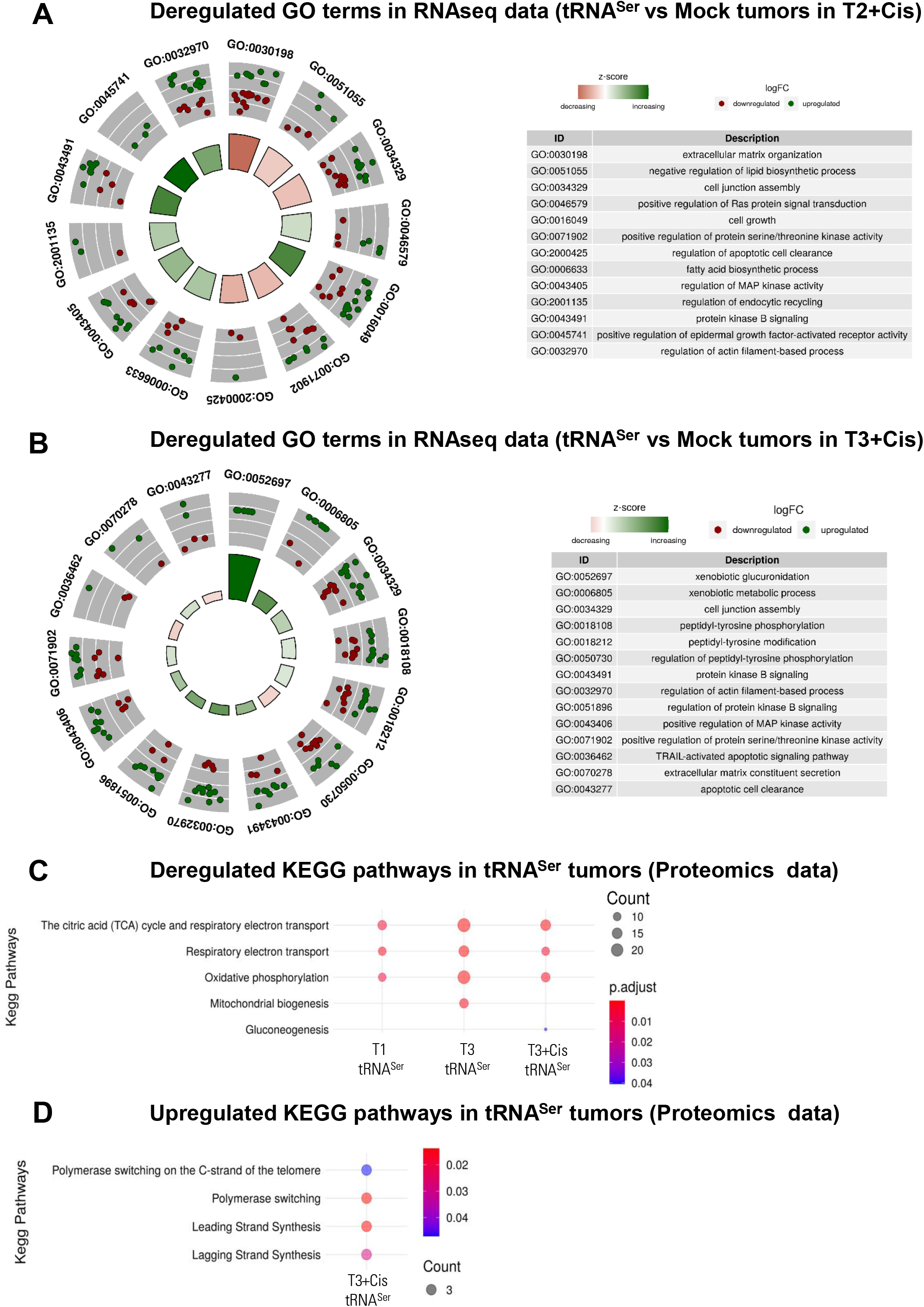
RNA-seq and proteomics confirm the activation of growth and survival pathways in tRNASer tumors. Circular plots depict deregulated GoTerms in tRNASer tumors. Color Scale: Red – downregulated; Green – upregulated. **A)** Deregulated GoTerms in tRNASer tumors in T2+Cis confirm the activation of pathways that are being affected by genomic variants found in this timepoint. **B)** Deregulated GoTerms in tRNASer tumors in T3+Cis confirm the activation of pathways that are being affected by genomic variants found in this timepoint. Overall, the upregulation of GO Terms confirms the activation of growth and survival signaling pathways, suggesting the presence of adaptive mutations in the genome. **C)** Deregulated KEGG pathways related with OXPHOS suppression in tRNASer tumors along the experiment found through proteomics analysis. **D)** Deregulated KEGG pathways in tRNASer tumors in T3+Cis retrieved from upregulated proteins at this timepoint showed evidence of polymerase switching and translesion synthesis.

Regarding the proteomics analysis, we could validate that “Mitochondrial Biogenesis” in T3+Cis tRNASer tumors is indeed affected (Fig. 3A). Interestingly OXPHOS related pathways are inactivated in T1 and T3 tRNASer tumors regardless of the treatment regimen, but not in T2 (Fig. 4C). Observing only the upregulated proteins in tRNASer tumors when compared to their Mock counterparts, we saw that these proteins are promoting polymerase switching and DNA synthesis in T3+Cis, which has been associated with Translesion synthesis (Fig. 4D) (32). This is a powerful mechanism that allow cells to thrive despite DNA damage (33). The decrease in OXPHOS and DNA damage repair are mechanisms that favor therapy resistance in NSCLC (34,35). Together, these results seem to point to a clonal selection in these tumors towards the acquisition of advantageous features that allow them to thrive under stressful conditions, such as the exposure to a cytotoxic agent.

### tRNASer tumors have a specific clonal evolution pattern

Tumors are a result of an evolutionary competition between subclones which display distinct somatic mutations that confer advantageous features (36). Since tRNASer tumors have an increasing number of variants with time and exposure to a trigger, we explored the clonal evolution of variants with “High”, “Moderate” and “Modifier” impacts using their allelic frequencies. As we have a huge number of variants we grouped the variants in 6 different groups: Group 1 – Variants present in Mock and tRNASer tumors throughout the whole experiment (921 variants); Group 2-Variants that are exclusive of tRNASer tumors and present throughout the whole experiment (100 variants); Group 3 – Exclusive variants of naïve/vehicle treated tRNASer tumors (27 variants); Group 4 – Exclusive variants of cisplatin treated tRNASer tumors (T2 and T3) (36 variants); Group 5 - Exclusive variants of cisplatin treated tRNASer tumors in T2 (130 variants); Group 6 – Exclusive variants of cisplatin treated tRNASer tumors in T3 (120 variants).

The vast majority of the cells (allelic frequency of the variants >0.7) display variants that were also present in Mock tumors (Group 1 variants) throughout the whole experiment. The allelic frequency of these variants varies just by 0.7% from T1 to T3+Cis (Fig. 5), indicating that these shared variants may be part of H460 identity as they are maintained in both Mock and tRNASer tumors independently of time and trigger exposure (Fig.5, Supl. Fig. 4). On the other hand, the allelic frequency of tRNASer exclusive variants (Group 2) increases by 3% from T1 to T3+Cis, suggesting they may be increasingly relevant for the tumor development along time and upon cisplatin treatment. Regarding the 286 variants appearing upon tRNASer tumor exposure to cisplatin (Groups 4, 5 and 6), their allelic frequency ranges from 0.25 to 0.29, meaning they are present in at least 25% of tumor cells. Given that these variants are activating growth and survival genes (Fig. 3, 4), this clonal adaptation driven by tRNASer upregulation is extremely relevant for tumor cell survival upon exposure to cisplatin.

**Figure 5.**
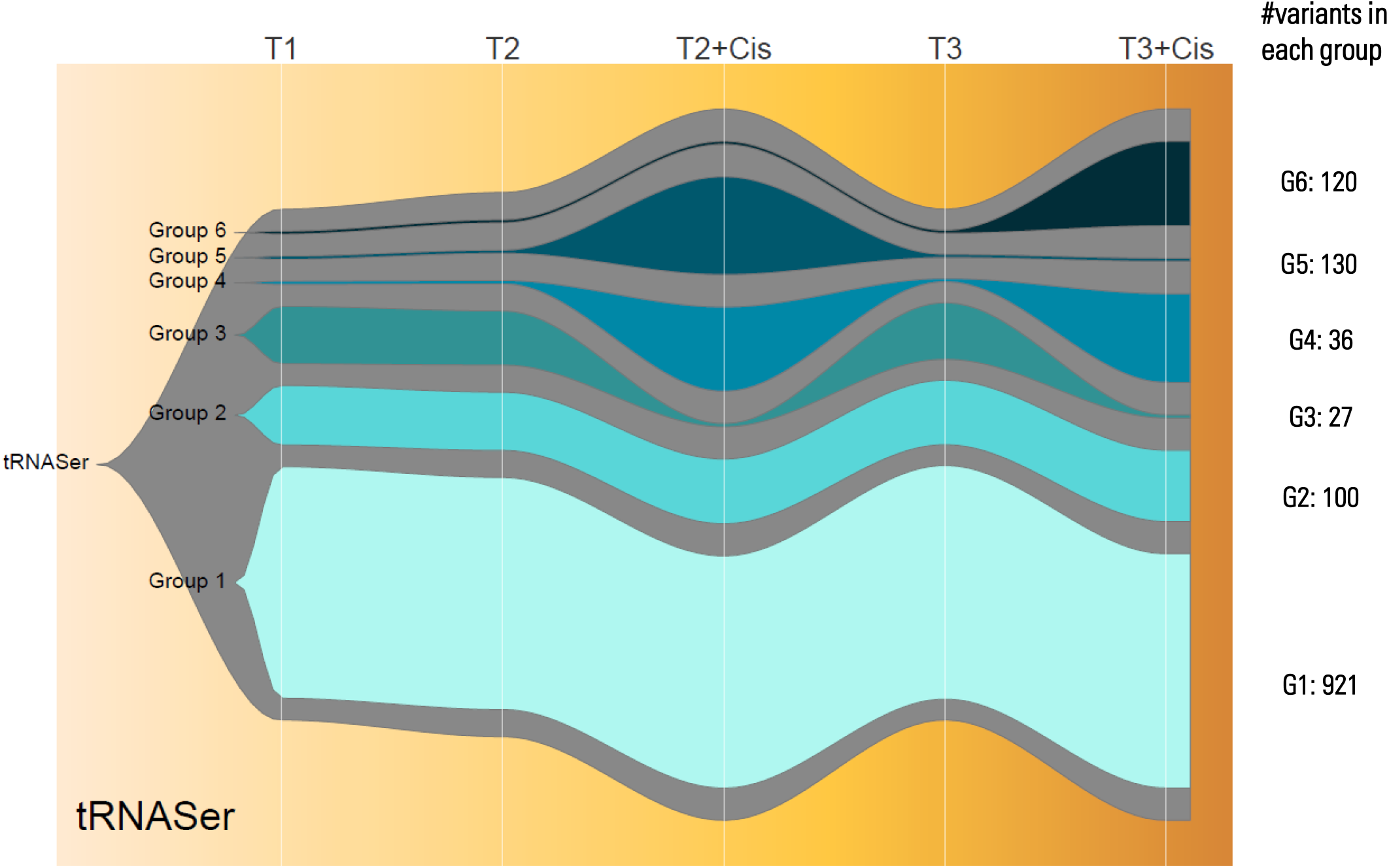
Clonal dynamics of genomic variants in tRNASer along time. Fish plot showing clonal dynamics of distinct groups of variants present along the experiment. The results suggest that clonal selection is important for tumor cell survival upon environmental stress, such as cisplatin exposure. Legend: Group 1 – Variants present in Mock and tRNASer tumors throughout the whole experiment; Group 2 – Variants exclusive of tRNASer tumors and present throughout the whole experiment; Group 3 – Exclusive variants of naïve/vehicle treated tRNASer tumors; Group 4 – Exclusive variants of cisplatin treated tRNASer tumors; Group 5 - Exclusive variants of cisplatin treated tRNASer tumors in T2; Group 6 – Exclusive variants of cisplatin treated tRNASer tumors in T3. The number of variants represented in each group is described in the right hand side of the graph.

## Discussion

Our work demonstrates for the first time that tRNASer deregulation increases the mutational load in cancer cells, activating signaling pathways that foster tumor cell survival and proliferation. Similar results were obtained in yeast, where the introduction of an ambiguous tRNASer induced adaptive mutations in the genome, rendering yeast more resistant to antifungals and environmental stressors (13,16). These results point to a cross-species mechanism to fixate important mutations driven by tRNA deregulation.

Our experiment was arguably too short to observe *de novo* mutations driven by exposure to cisplatin in 3 or more tumors. Therefore, the most probable explanation for the mutations that arise with time and cisplatin exposure is the selection of the fittest cell population. In this case, all the variants observed were already present in the cells that were inoculated in mice but in small proportions that were undetectable in T1. With time and cisplatin exposure those variants became more relevant for tumor cell survival and took over the previous cell population. This is a well-known phenomenon in tumors called clonal evolution (37). Tumors are not constituted by a static and uniform cell population, but rather by a dynamic and heterogeneous mixture of different cell types and several cell subclones (38,39). These clones are the primary reservoir of intratumoral genetic diversity, fostering tumor adaptation to unfavorable conditions (40). On the other hand, the deregulation of tRNAs upon environmental cues is common, not only in cancer, but also in other pathologies (41,42). Their deregulation was already implicated in several non-canonical functions from inhibition of cell death (43) to increased cell metabolism (44). Our work expands these functions in mammalian cells, by showing that tumors with tRNASer deregulation have an increased number of variants, especially synonymous and missense variants (Fig. 1A, 2). The analysis performed show a strong correlation among results, as these tumors also present high TMB and hypermutator and MMR deficiency signatures (Fig. 6).

**Figure 6.**
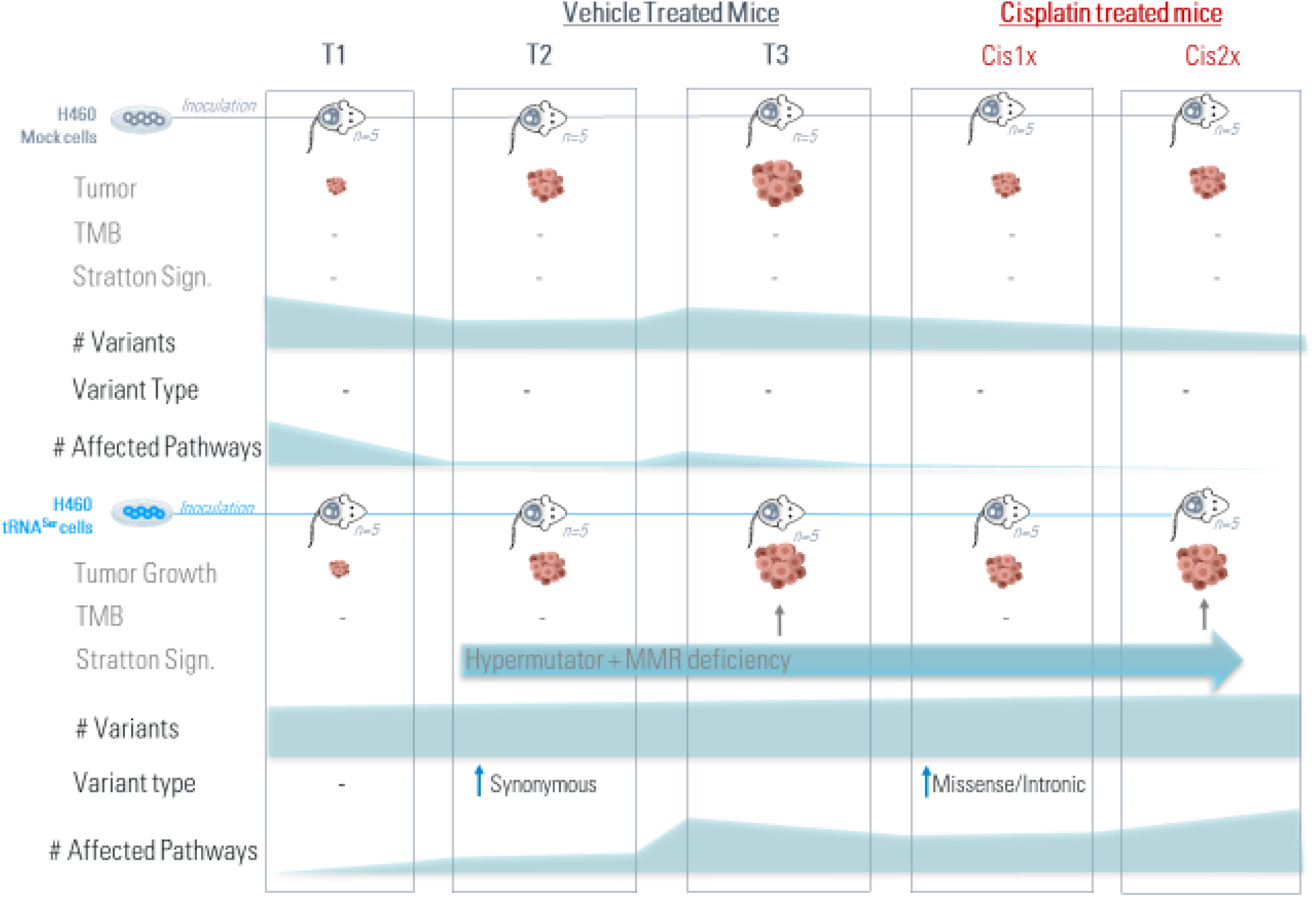
tRNASer is a new driver of genomic instability with important adaptive consequences. To summarize all our work, we observed that overexpression of tRNASer, which is present in tumors with bad prognosis, renders the tumors insensitive to cisplatin treatment, increases the tumor mutation burden upon treatment, increases Hypermutator and MMR-deficiency signatures, and increases the number of variants that affect predominantly growth and survival genes. This means that the deregulation of a single tRNA is enough to fixate important and adaptive mutations in cancer cells, highlighting a new and unexplored mechanism of cancer cells.

From the TCGA data analyzed in tRIC database (45), we can observe that there are tumors with high and low expression of tRNASer (Fig. S3). Moreover, tumors with increased tRNASer expression have a worse overall survival (Cox *pvalue* = 0.00029), which correlates nicely with our results, since we observed that tRNASer tumors do not respond to cisplatin treatment (Fig. S2). This particular tRNASer has also been associated with worse 5-year overall survival and risk of recurrence in breast cancer (24), suggesting these results may be transversal to several types of cancer. We disclosed that the variants present in tRNASer tumors were affecting genes related with cell survival and growth pathways (Fig. 3). The RNA-seq and proteomics further confirmed the activation of these pathways (Fig. 4). Interestingly, we observed an inhibition of OXPHOS and it has been documented that tRNAs can bind cytochrome C and inhibit mitochondria function (43), shifting cell metabolism to anaerobic respiration (46). These adaptive mutations prime tRNASer tumors for plasticity, conferring them an intrinsic capability to thrive in stressful environments, such as exposure to a cytotoxic agent (47,48).

Therefore, our work unveils an additional mechanism by which tumor cells can increase their genetic diversity reservoir with important functional consequences for tumor development and possibly therapy resistance.

## Conclusion

The deregulation of tRNAs in tumor cells promotes genomic instability, increasing the tumor mutational burden and fixating mutations that result in increased fitness and resistance to environmental stresses, such as chemotherapy.

## Methodology

### Mice Experiments

Six-week-old male N:NIH(s)II:nu/nu nude mice were obtained previously from the Medical School, University of Cape Town in 1991 and then reproduced, maintained and housed at IPATIMUP Animal House at the Medical Faculty of the University of Porto, in a pathogen-free environment under controlled conditions of light and humidity. Animal experiments were conducted at the i3S Animal House in accordance with the Guidelines for the Care and Use of Laboratory Animals, directive 2010/63/EU and the ethical approval of DGAV (reference no. 0421/000/000/2017). Male nude mice, aged 6-8 weeks, were used for *in vivo* experiments. H460 cell lines harboring the empty vector (Mock) or the tRNA^Ser^(AGA) were subcutaneously injected in the dorsal flanks using a 25-gauge needle (1×10^6^ /mouse). A total of 5 mice per group were used according to the scheme in Fig. S1. Mice were weighed, and tumor width and length were measured with calipers three times per week. Tumor volumes were calculated assuming ellipsoid growth patterns. T1 mice were humanely euthanized when tumors reached ∼250mm^3^, T2 mice were euthanized when tumors reached a volume of 1500mm^3^ and T3 tumors were euthanized when tumors reached ∼1800mm^3^. Tumors were collected, fixed in 10% buffered formalin, paraffin embedded and then sectioned for histopathological examination. A part of each tumor was frozen in liquid nitrogen for DNA, RNA and protein extraction and stored at –80ºC until used.

### WES (including sample preparation)

Frozen tumor pieces were homogenized using liquid nitrogen. gDNA was extracted from 25mg of frozen tissue using the GenElute™ Mammalian Genomic DNA Miniprep kit (G1N70-1KT, Merck). gDNA quantity was measured using a NanoDrop Spectophotometer and its integrity was assessed by running 100ng of DNA in a 1% agarose gel. 2μg of DNA were sent to Macrogen to be sequenced. Libraries for WES were constructed using the Agilent SureSelect XT_V6 library and were sequenced on a NovaSeq Illumina sequencer with a 200x coverage and a read length of 150bp paired-end.

Initial quality checking of the raw reads was performed with FASTQC v0.11.7 (Andrews S, 2010) and parameters were adjusted to improve the quality of the reads with Trimmomatic v0.36 (Bolger et al, 2014). The reads were mapped against the Human reference genome (hg38) using BWA-MEM v0.7.7 with default options.

### Variant analysis

Variant calling were made with GATK v4.1.1.3, using best practices. At the end we filtered all variants to match the amplified regions made by SureSelect v7 hg38 kit (Agilent), and were annotated with Variant Effect Prediction (VEP) release 104.

### Tumor mutation Burden

The Tumor Mutation Burden (TMB) was calculated for each sample according to the following formula:

*#non-synonymous variants/ 49*.*8 Mb*

### Stratton Signatures

Mutation signatures from MAF’s were obtained with scripts available in (https://github.com/mskcc/mutation-signatures v3.2)

### Clonal Evolution Analysis and Fish plot

Clonal evolution was assessed based on the allelic frequencies of variants with “High”, “Moderate” and “Modifier” impacts for each condition and time point.

The Allelic Frequencies for each variant were computed by evaluating the relative abundance of different alleles within a sample, taking into account the average across all replicates of that sample to determine the final value. Only variants that were observed in at least three out of five replicates were taken into consideration.

To obtain the final value, all the Allele Frequencies averages, for each variant, where computed again, making a new global average for each time point, taking in consideration impacts and six categories that correspond to the following groups: i) Group 1 – Variants present in Mock and tRNASer tumors throughout the whole experiment (921 variants); ii) Group 2-Variants that are exclusive of tRNASer tumors and present throughout the whole experiment (100 variants); iii) Group 3 – Exclusive variants of naïve/vehicle treated tRNASer tumors (27 variants); iv) Group 4 – Exclusive variants of cisplatin treated tRNASer tumors (T2 and T3) (36 variants); v) Group 5 - Exclusive variants of cisplatin treated tRNASer tumors in T2 (130 variants); vi) Group 6 –Exclusive variants of cisplatin treated tRNASer tumors in T3 (120 variants).

This global Allele Frequency average, by time points, were then used to create the fish plots, based on the fishplot R package (https://github.com/chrisamiller/fishplot), and the code was made available on the github (https://github.com/ibigen/fishplot_tRNA_MOCK).

### RNAseq

RNA for Whole Transcriptome Sequencing (RNAseq) was extracted from 3 tumors from each timepoint and derived-cell line using the mirVana miRNA Isolation Kit (Invitrogen, Oregon, USA), according to the kit’s instruction manual and maintained at - 80ºC until further processing. RNA quality analysed using Agilent2100-Bioanalyzer (RIN>8). 8ug of total RNA were sent to Macrogen to be sequenced. Libraries for RNAseq were constructed using Illumina TruSeq stranded mRNA library kit. 30M reads were obtained for each sample with a read length of 100bp paired-end. Unique-reads were mapped to hg38 human genome using Bowtie, TopHat2 and differentially-expressed genes (DEGs) detected using *DEseq2* R package. Genes with log2fold-change>1 or <-1 and corrected p<0.01, were considered DEGs. Statistics performed using R. *ClusterProfiler* R package was used for assessment of significantly-enriched GO terms and pathways (*padj*<0.05).

### Proteomics

25 mg of tumor tissue (3 for each timepoint) were homogenized in Protein Lysis Buffer (0.5% Triton X-100, 50mM HEPES, 250mM NaCl, 1mM DTT, 1mM NaF, 2mM EDTA, 1mM EGTA, 1mM PMSF, 1mM Na3VO4 supplemented with a cocktail of protease inhibitors (Complete, EDTA-free, Roche). Tissue was sonicated with a probe sonicator in 5 pulses of 5 seconds, incubated on ice for 30min and centrifuged at 20000*g* for 30min at 4ºC. 10 mL of the supernatant were used to measure protein concentration with BCA assay (Thermo Fisher Scientific). 100 mg of total protein were sent to the i3S proteomics facility. After reduction, alquylation and tryptic digestion, the resulting peptides were quantified. 500ng of each sample were injected in the mass spectrometer for analysis. Protein identification was done using the human reference proteome retrieved from UniProt and a database of mass spectra. We followed the Label Free Quantification Method tacking into account the pre-designed timepoints of the mice experiment using the Proteome Discoverer software.

## Availability of data and materials

The cell lines used analyzed during the current study are available from the corresponding author on reasonable request. The datasets used and/or analyzed during the current study will be made available upon publication. Proteomics data is deposited in iProX repository (ID: IPX0006267000), WES and RNAseq will be made available upon reasonable request to the corresponding author.

## Author Contributions

Conceptualization, M.S., M.A.S.S. and C.O..; methodology, M.S., M.F., M.P., A.R., S.R and A.A.; software, M.P., A.R. and M.F.; validation, M.S., M.F. and C.O.; formal analysis, M.S., M.F., M.P., A.R. and C.O.; investigation, M.S., M.F., M.P., A.R. and C.O.; resources, M.A.S.S., M.S. and C.O.; data curation, M.S., M.F., M.P., A.R. and C.O. ; writing— M.S. and C.O.; writing—review and editing, M.S. and C.O; visualization, M.S., M.F., M.P., A.R. and C.O. ; supervision, C.O.; project administration, M.S. and C.O.; funding acquisition, M.S. and C.O. All authors have read and agreed to the published version of the manuscript.

## Acknowledgements

The authors acknowledge the efforts of the Scientific Platforms staff at i3S, namely Hugo Osório from the Proteomics facility, Ana Mafalda Rocha from the Genomics facility and the veterinarians and caretakers from the Animal Facility, who were essential for the outcome of this project.

## References

1. Hanahan D, Weinberg RA. Hallmarks of cancer: The next generation. Vol. 144, Cell. 2011. p. 646–74.

2. Langie SAS, Koppen G, Desaulniers D, Al-Mulla F, Al-Temaimi R, Amedei A, et al. Causes of genome instability: the effect of low dose chemical exposures in modern society. Carcinogenesis [Internet]. 2015 Jun 1;36(Suppl_1):S61–88. Available from: https://doi.org/10.1093/carcin/bgv031

3. Sonugür FG, Akbulut H. The Role of Tumor Microenvironment in Genomic Instability of Malignant Tumors [Internet]. Vol. 10, Frontiers in Genetics. 2019. p. 1063. Available from: https://www.frontiersin.org/article/10.3389/fgene.2019.01063

4. Takeshima H, Ushijima T. Accumulation of genetic and epigenetic alterations in normal cells and cancer risk. npj Precis Oncol [Internet]. 2019;3(1):7. Available from: https://doi.org/10.1038/s41698-019-0079-0

5. Minakawa Y, Shimizu A, Matsuno Y, Yoshioka K-I. Genomic Destabilization Triggered by Replication Stress during Senescence. Cancers (Basel). 2017 Nov 21;9:159.

6. Basu AK. DNA Damage, Mutagenesis and Cancer. Int J Mol Sci [Internet]. 2018 Mar 23;19(4):970. Available from: https://pubmed.ncbi.nlm.nih.gov/29570697

7. Burgess JT, Rose M, Boucher D, Plowman J, Molloy C, Fisher M, et al. The Therapeutic Potential of DNA Damage Repair Pathways and Genomic Stability in Lung Cancer. Front Oncol [Internet]. 2020 Jul 28;10:1256. Available from: https://pubmed.ncbi.nlm.nih.gov/32850380

8. Fan Y, Zhu X, Xu Y, Lu X, Xu Y, Wang M, et al. Cell-Cycle and DNA-Damage Response Pathway Is Involved in Leptomeningeal Metastasis of Non–Small Cell Lung Cancer. Clin Cancer Res [Internet]. 2018 Jan 1;24(1):209 LP –216. Available from: http://clincancerres.aacrjournals.org/content/24/1/209.abstract

9. Bakhoum SF, Cantley LC. The Multifaceted Role of Chromosomal Instability in Cancer and Its Microenvironment. Cell [Internet]. 2018 Sep 6;174(6):1347–60. Available from: https://pubmed.ncbi.nlm.nih.gov/30193109

10. Pua KH, Chew CL, Lane DP, Tergaonkar V. Inflammation-associated genomic instability in cancer. Genome Instab Dis [Internet]. 2020;1(1):1–9. Available from: https://doi.org/10.1007/s42764-019-00006-6

11. Sophie T, Pauline F, Laetitia L, Deborah G, Patricia M, Matteo S, et al. The Genotoxin Colibactin Shapes Gut Microbiota in Mice. mSphere [Internet]. 2020 Jul 1;5(4):e00589–20. Available from: https://doi.org/10.1128/mSphere.00589-20

12. Cao Y, Oh J, Xue M, Huh WJ, Wang J, Gonzalez-Hernandez JA, et al. Commensal microbiota from patients with inflammatory bowel disease produce genotoxic metabolites. Science (80-). 2022;378(6618).

13. Bezerra AR, Simões J, Lee W, Rung J, Weil T, Gut IG, et al. Reversion of a fungal genetic code alteration links proteome instability with genomic and phenotypic diversification. Proc Natl Acad Sci U S A [Internet]. 2013;110:11079–84. Available from: http://www.pubmedcentral.nih.gov/articlerender.fcgi?artid=3704024&tool=pmcentrez&rendertype=abstract

14. Lee JY, Kim DG, Kim B-G, Yang WS, Hong J, Kang T, et al. Promiscuous methionyl-tRNA synthetase mediates adaptive mistranslation to protect cells against oxidative stress. J Cell Sci [Internet]. 2014 Sep 30;127(19):4234 LP –4245. Available from: http://jcs.biologists.org/content/127/19/4234.abstract

15. Kalapis D, Bezerra AR, Farkas Z, Horvath P, Bódi Z, Daraba A, et al. Evolution of Robustness to Protein Mistranslation by Accelerated Protein Turnover. PLoS Biol. 2015;13(11).

16. Weil T, Santamaría R, Lee W, Rung J, Tocci N, Abbey D, et al. Adaptive Mistranslation Accelerates the Evolution of Fluconazole Resistance and Induces Major Genomic and Gene Expression Alterations in <em>Candida albicans</em>. Mitchell AP, editor. mSphere [Internet]. 2017 Aug 9;2(4). Available from: http://msphere.asm.org/content/2/4/e00167-17.abstract

17. Guimarães AR, Correia I, Sousa I, Oliveira C, Moura G, Bezerra AR, et al. tRNAs as a Driving Force of Genome Evolution in Yeast. Vol. 12, Frontiers in Microbiology. Frontiers Media S.A.; 2021.

18. Raab JR, Chiu J, Zhu J, Katzman S, Kurukuti S, Wade PA, et al. Human tRNA genes function as chromatin insulators. EMBO J [Internet]. 2012;31(2):330–50. Available from: http://www.pubmedcentral.nih.gov/articlerender.fcgi?artid=3261562&tool=pmcentrez&rendertype=abstract

19. Donze D, Kamakaka RT. RNA polymerase III and RNA polymerase II promoter complexes are heterochromatin barriers in Saccharomyces cerevisiae. EMBO J [Internet]. 2001 Feb 1;20(3):520–31. Available from: https://doi.org/10.1093/emboj/20.3.520

20. Ebersole T, Kim JH, Samoshkin A, Kouprina N, Pavlicek A, White RJ, et al. tRNA genes protect a reporter gene from epigenetic silencing in mouse cells. Cell Cycle. 2011 Aug 15;10(16):2779–91.

21. Iwasaki Y, Ikemura T, Kurokawa K, Okada N. Implication of a new function of human tDNAs in chromatin organization. Sci Rep. 2020 Dec 1;10(1).

22. Goodarzi H, Nguyen HCB, Zhang S, Dill BD, Molina H, Tavazoie SF. Modulated Expression of Specific tRNAs Drives Gene Expression and Cancer Progression. Cell. 2016;(165):1416–1427.

23. Santos M, Pereira PM, Varanda AS, Carvalho J, Azevedo M, Mateus DD, et al. Codon misreading tRNAs promote tumor growth in mice. RNA Biology. 2018;1–14.

24. Krishnan P, Sunita G, Wang B, Heyns M, Li D, Mackey JR, et al. Genome-wide profiling of transfer RNAs and their role as novel prognostic markers for breast cancer. Sci Rep. 2016;6(32843).

25. Santos M, Fidalgo A, Varanda AS, Soares AR, Almeida GM, Martins D, et al. Upregulation of tRNA-Ser-AGA-2-1 Promotes Malignant Behavior in Normal Bronchial Cells. Front Mol Biosci. 2022;9.

26. Gingold H, Tehler D, Christoffersen NR, Nielsen MM, Asmar F, Kooistra SM, et al. A Dual Program for Translation Regulation in Cellular Proliferation and Differentiation. Cell [Internet]. 2015 Jun 20;158(6):1281–92. Available from: http://dx.doi.org/10.1016/j.cell.2014.08.011

27. Xu Z, Dai J, Wang D, Lu H, Dai H, Ye H, et al. Assessment of tumor mutation burden calculation from gene panel sequencing data. Onco Targets Ther [Internet]. 2019 May 6;12:3401–9. Available from: https://pubmed.ncbi.nlm.nih.gov/31123404

28. Willis C, Fiander M, Tran D, Korytowsky B, Thomas J-M, Calderon F, et al. Tumor mutational burden in lung cancer: a systematic literature review. Oncotarget [Internet]. 2019 Nov 12;10(61):6604–22. Available from: https://pubmed.ncbi.nlm.nih.gov/31762941

29. Strickler JH, Hanks BA, Khasraw M. Tumor Mutational Burden as a Predictor of Immunotherapy Response: Is More Always Better? Clin Cancer Res [Internet]. 2021 Mar 1;27(5):1236 LP –1241. Available from: http://clincancerres.aacrjournals.org/content/27/5/1236.abstract

30. Alexandrov LB, Kim J, Haradhvala NJ, Huang MN, Tian Ng AW, Wu Y, et al. The repertoire of mutational signatures in human cancer. Nature [Internet]. 2020;578(7793):94–101. Available from: https://doi.org/10.1038/s41586-020-1943-3

31. Leite M, Corso G, Sousa S, Milanezi F, Afonso LP, Henrique R, et al. MSI phenotype and MMR alterations in familial and sporadic gastric cancer. Int J cancer. 2011 Apr;128(7):1606–13.

32. Zhao L, Washington MT. Translesion Synthesis: Insights into the Selection and Switching of DNA Polymerases. Vol. 8, Genes. 2017.

33. Lange SS, Takata K, Wood RD. DNA polymerases and cancer. Nat Rev Cancer [Internet]. 2011 Feb;11(2):96–110. Available from: https://pubmed.ncbi.nlm.nih.gov/21258395

34. Ashton TM, McKenna WG, Kunz-Schughart LA, Higgins GS. Oxidative Phosphorylation as an Emerging Target in Cancer Therapy. Clin Cancer Res [Internet]. 2018 Jun 1;24(11):2482 LP –2490. Available from: http://clincancerres.aacrjournals.org/content/24/11/2482.abstract

35. Liang X, Qin Q, Liu B, Li X, Zeng L, Wang J, et al. Targeting DNA-PK overcomes acquired resistance to third-generation EGFR-TKI osimertinib in non-small-cell lung cancer. Acta Pharmacol Sin [Internet]. 2021;42(4):648–54. Available from: https://doi.org/10.1038/s41401-020-00577-1

36. Rampias T. Exploring the Eco-Evolutionary Dynamics of Tumor Subclones. Vol. 12, Cancers. 2020.

37. Greaves M, Maley CC. Clonal evolution in cancer. Nature [Internet]. 2012;481(7381):306–13. Available from: https://doi.org/10.1038/nature10762

38. Shlush LI, Hershkovitz D. Clonal Evolution Models of Tumor Heterogeneity. Am Soc Clin Oncol Educ B [Internet]. 2015 May 1;(35):e662–5. Available from: https://doi.org/10.14694/EdBook_AM.2015.35.e662

39. Altorki NK, Markowitz GJ, Gao D, Port JL, Saxena A, Stiles B, et al. The lung microenvironment: an important regulator of tumour growth and metastasis. Nat Rev Cancer [Internet]. 2019;19(1):9–31. Available from: https://doi.org/10.1038/s41568-018-0081-9

40. Losic B, Craig AJ, Villacorta-Martin C, Martins-Filho SN, Akers N, Chen X, et al. Intratumoral heterogeneity and clonal evolution in liver cancer. Nat Commun [Internet]. 2020;11(1):291. Available from: https://doi.org/10.1038/s41467-019-14050-z

41. Santos M, Fidalgo A, Varanda AS, Oliveira C, Santos MAS. tRNA Deregulation and Its Consequences in Cancer. Trends Mol Med [Internet]. 2019 Oct 1;25(10):853–65. Available from: https://doi.org/10.1016/j.molmed.2019.05.011

42. Avcilar-Kucukgoze I, Kashina A. Hijacking tRNAs From Translation: Regulatory Functions of tRNAs in Mammalian Cell Physiology [Internet]. Vol. 7, Frontiers in Molecular Biosciences. 2020. Available from: https://www.frontiersin.org/article/10.3389/fmolb.2020.610617

43. Mei Y, Yong J, Liu H, Shi Y, Meinkoth J, Dreyfuss G, et al. tRNA Binds to Cytochrome c and Inhibits Caspase Activation. Mol Cell [Internet]. 2010;37(5):668–78. Available from: http://www.sciencedirect.com/science/article/pii/S1097276510000742

44. Pavon-Eternod M, Gomes S, Rosner MR, Pan T. Overexpression of initiator methionine tRNA leads to global reprogramming of tRNA expression and increased proliferation in human epithelial cells. RNA [Internet]. 2013;19:461–6. Available from: http://www.ncbi.nlm.nih.gov/pubmed/23431330

45. Zhang Z, Ruan H, Liu C-J, Ye Y, Gong J, Diao L, et al. tRic: a user-friendly data portal to explore the expression landscape of tRNAs in human cancers. RNA Biol. 2020 Nov;17(11):1674–9.

46. Dhakal R, Tong C, Anderson S, Kashina AS, Cooperman B, Bau HH. Dynamics of intracellular stress-induced tRNA trafficking. Nucleic Acids Res [Internet]. 2019 Feb 28;47(4):2002–10. Available from: https://doi.org/10.1093/nar/gky1208

47. Menchón SA. The Effect of Intrinsic and Acquired Resistances on Chemotherapy Effectiveness. Acta Biotheor [Internet]. 2015;63(2):113–27. Available from: http://dx.doi.org/10.1007/s10441-015-9248-x

48. Adachi Y, Ito K, Hayashi Y, Kimura R, Tan TZ, Yamaguchi R, et al. Epithelial-to-Mesenchymal Transition is a Cause of Both Intrinsic and Acquired Resistance to KRAS G12C Inhibitor in KRAS G12C–Mutant Non–Small Cell Lung Cancer. Clin Cancer Res [Internet]. 2020 Nov 15;26(22):5962 LP –5973. Available from: http://clincancerres.aacrjournals.org/content/26/22/5962.abstract

