## Supplementary figures and images for "tRNA^Ser^ overexpression induces adaptive mutations in NSCLC tumors"

### Suplemmentary Figures 1-4

#### Supplementary Figure 1

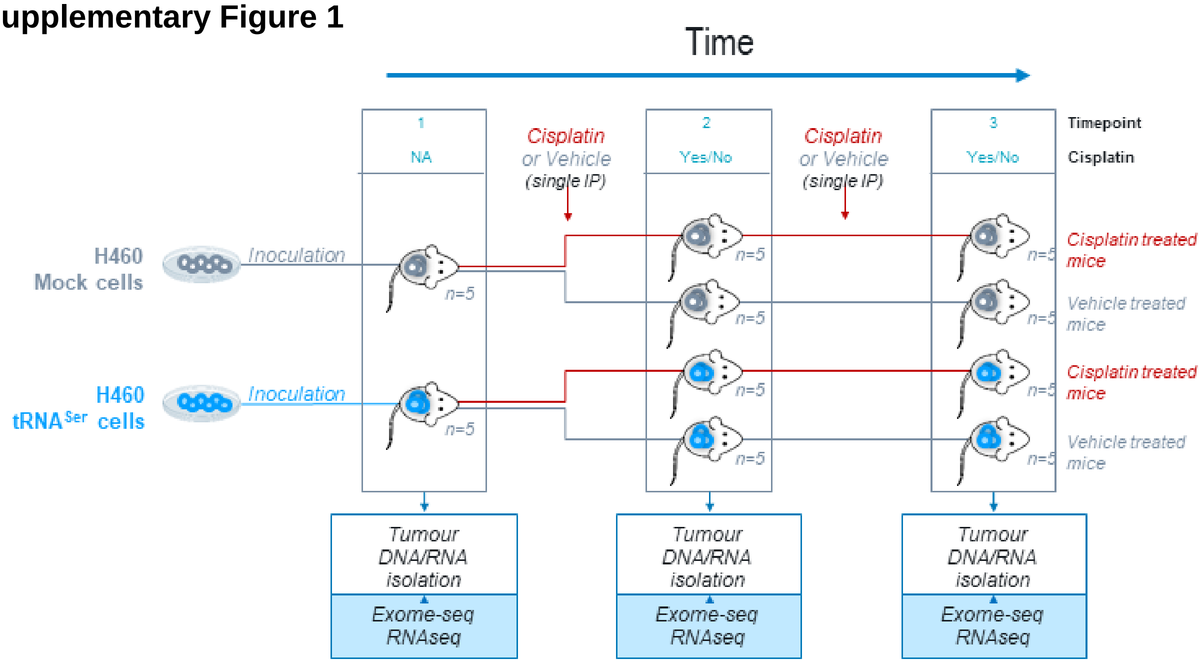


#### Supplementary Figure 2

**A**

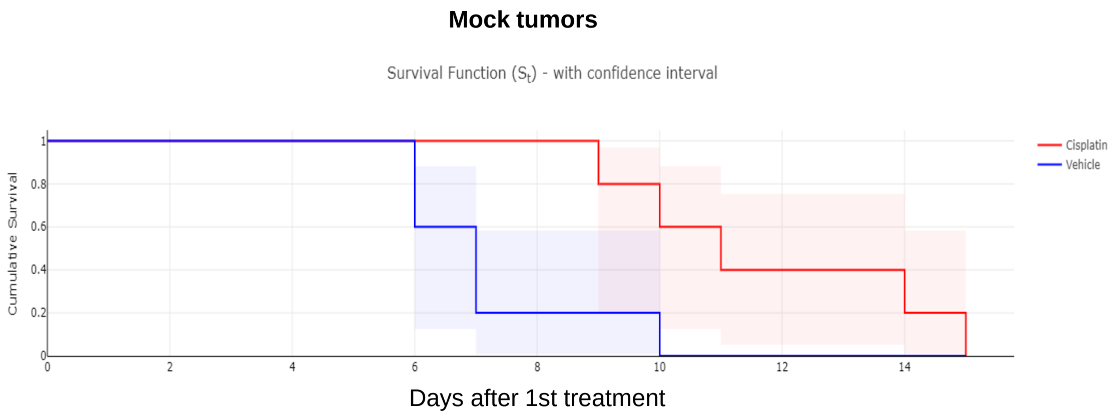


**B**

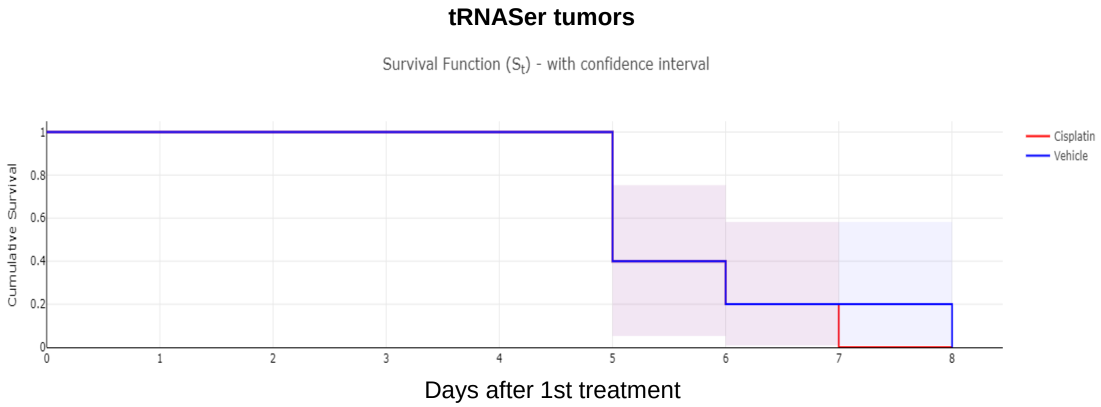

#### Supplementary Figure 3

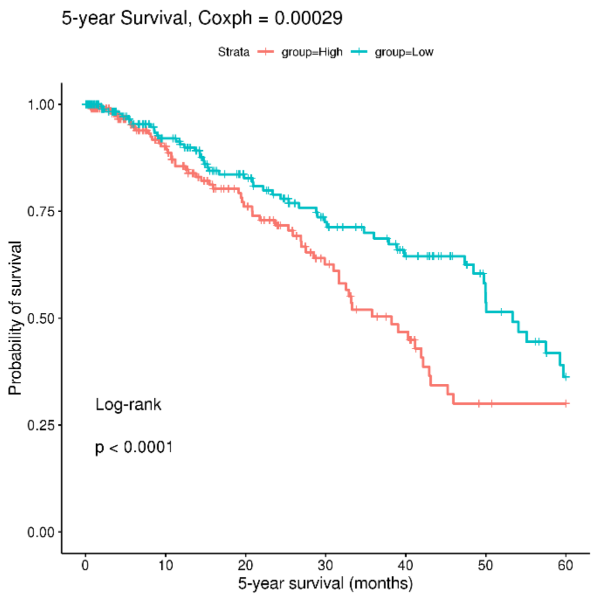


#### Supplementary Figure 4

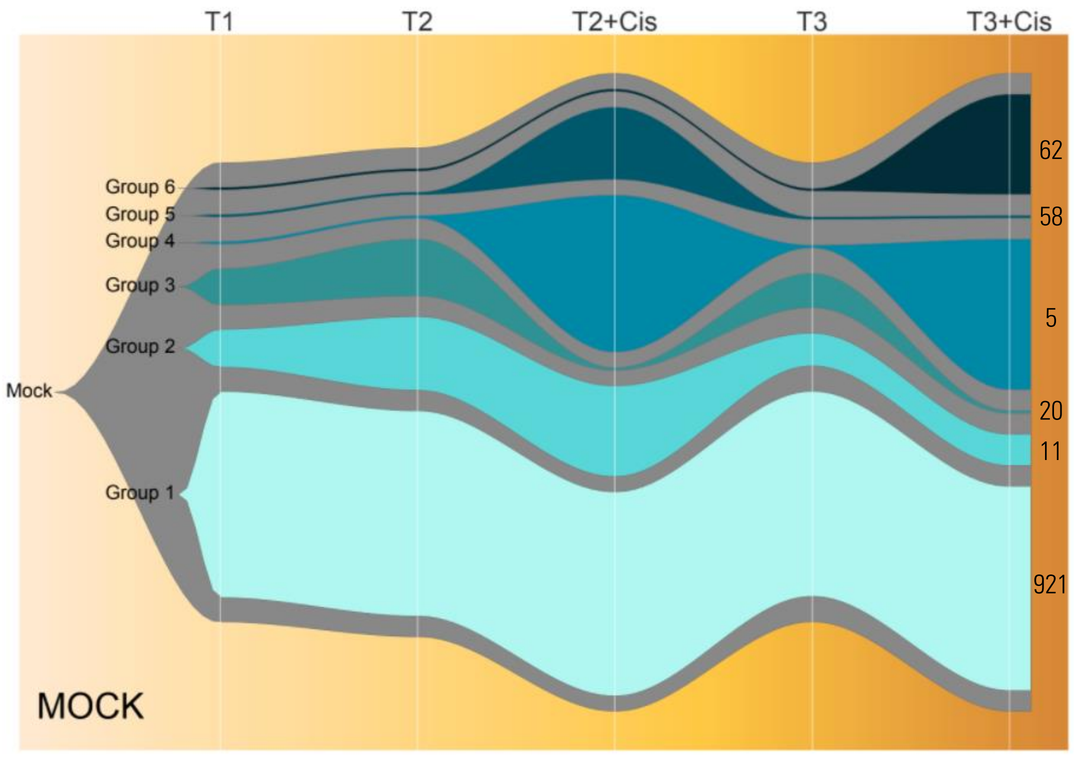
